# Learning alters neural activity to simultaneously support memory and action

**DOI:** 10.1101/2022.07.05.498856

**Authors:** Darby M. Losey, Jay A. Hennig, Emily R. Oby, Matthew D. Golub, Patrick T. Sadtler, Kristin M. Quick, Stephen I. Ryu, Elizabeth C. Tyler-Kabara, Aaron P. Batista, Byron M. Yu, Steven M. Chase

## Abstract

How are we able to learn new behaviors without disrupting previously learned ones? To understand how the brain achieves this, we used a brain-computer interface (BCI) learning paradigm, which enables us to detect the presence of a memory of one behavior while performing another. We found that learning to use a new BCI map altered the neural activity that monkeys produced when they returned to using a familiar BCI map, in a way that was specific to the learning experience. That is, learning left a “memory trace.” This memory trace co-existed with proficient performance under the familiar map, primarily by altering dimensions of neural activity that did not impact behavior. Such a memory trace could provide the neural underpinning for the joint learning of multiple motor behaviors without interference.

## Introduction

Suppose an experienced skier learns to snowboard. Skiing and snowboarding require different sets of muscle activations, driven by different neural population activity patterns, to achieve the same goal of getting down the mountain without falling. How does the brain incorporate the knowledge about how to snowboard along with the knowledge about how to ski? The first possibility is that after learning to snowboard the neural activity used for skiing remains unchanged. In this scenario, the new neural activity for snowboarding would only be recalled when snowboarding again. Such context-dependent recall has been observed in certain learning settings, such as the remapping of hippocampal place-fields between environments (Alme et al., 2014), and has been proposed as a potential mechanism for motor memory storage (Herzfeld et al., 2014).

The second possibility is that the neural activity used for skiing is altered by the recently acquired ability to snowboard. Several studies have suggested that this might be the case, as neural tuning has been observed to change as a result of motor adaptation (Li et al., 2001, Arce et al., 2010, Cherian et al., 2013, Perich and Miller, 2017, Sun et al., 2022). There are two possible reasons for these neural activity changes. First, these changes might constitute a memory of the learning experience. That is, learning could lead to a “memory trace”, which we define as an alteration of the population activity patterns used to perform familiar tasks in a manner that renders them also appropriate for a newly learned task. Second, these changes could be attributed to the many task-agnostic factors, such as changes in arousal (Cowley et al., 2020, Hennig et al., 2021a), motivation (Roesch and Olson, 2004), posture (Graziano, 2006), or altered arm dynamics (Cherian et al., 2013, Perich and Miller, 2017). Without a known causal link between neural activity and behavior, it is difficult to determine if and how changes in neural activity after learning might constitute a memory trace.

Here we overcome this difficulty by leveraging a brain-computer interface (BCI) paradigm (Jarosiewicz et al., 2008, Ganguly and Carmena, 2009, Koralek et al., 2012, Hwang et al., 2013, Sadtler et al., 2014, Gulati et al., 2017, Jeon et al., 2022, Oby et al., 2019). A key advantage of a BCI for the study of motor memory is that the relationship between neural activity and behavior (termed the BCI map) is specified by the experimenter (Golub et al., 2016, Orsborn and Pesaran, 2017). This feature of a BCI is crucial in enabling us to look for a memory trace because it allows us to evaluate how changes in neural activity relate to a task that is not being performed.

We trained three monkeys to perform a BCI task. We used two different BCI maps in each experimental session. Much like the example of an experienced skier learning to snowboard, a monkey first controlled a computer cursor using a familiar *Map A*, and then learned how to use a new *Map B*. Following learning, we reinstated Map A. This allowed us to evaluate whether monkeys used different population activity patterns to control Map A before and after learning Map B. Furthermore, to see if neural activity showed a memory trace of having learned Map B, we evaluated how well the neural activity produced by the monkey during the use of Map A would have controlled the cursor through the offline Map B, comparing pre-versus post-learning. We observed that, after learning Map B, the monkeys were subsequently able to control the cursor using Map A, and yet the neural activity remained consistent with improved performance using Map B. That is, learning left a memory trace by altering the neural activity used to perform the familiar task, such that the neural activity became more appropriate for the learned task. The memory trace coexisted alongside proficient Map A performance by altering neural activity primarily along dimensions that did not affect cursor movements under Map A. Overall, our results reveal that learning can leave a memory trace in neural population activity that need not interfere with the subsequent behavior. The formation of a memory trace may thus provide a mechanism to facilitate the learning of multiple motor skills without interference (Krakauer et al., 2005), instantaneous switching between tasks, and rapid relearning of motor behaviors (“savings”).

## Results

Here we study how learning to perform a new task affects the neural activity used while performing a familiar task (Fig. 1a). We trained three monkeys to perform an eight-target center-out task using a brain-computer interface (BCI). The mon-key’s goal on each trial was to guide a computer cursor to an instructed target by modulating his neural activity (Fig. 1b; see Methods). At each moment in time, a BCI map determined the relationship between the neural activity, recorded from ∼90 neural units in primary motor cortex (M1), and the cursor’s 2D velocity. Each experiment utilized two different BCI maps, *Map A* and *Map B*, presented across three blocks of trials (Fig. 1c). During the first block (“Task A1”), we provided the monkeys with Map A, which was an “intuitive” map calibrated that day to allow for proficient cursor control without any learning. For the second block (“Task B”), we changed the BCI map to Map B, which the monkey had never used before (see Methods). This resulted in an initial decrement in the monkey’s performance, which improved over the course of several hundred trials as he learned to control the cursor. In the third block (“Task A2”), we reinstated Map A. This typically resulted in the well-known aftereffect that typically follows a bout of motor learning after which performance returns to level comparable to that of Task A1 (Shadmehr and Mussa-Ivaldi, 1994). Data from the Task A1 and Task B periods have been examined in prior work (Sadtler et al., 2014, Golub et al., 2018, Hennig et al., 2018, 2021a). In this study, we now focus on the neural activity recorded during Task A2, which is the appropriate epoch to address our central question and which we have not reported on previously.

**Figure 1.**
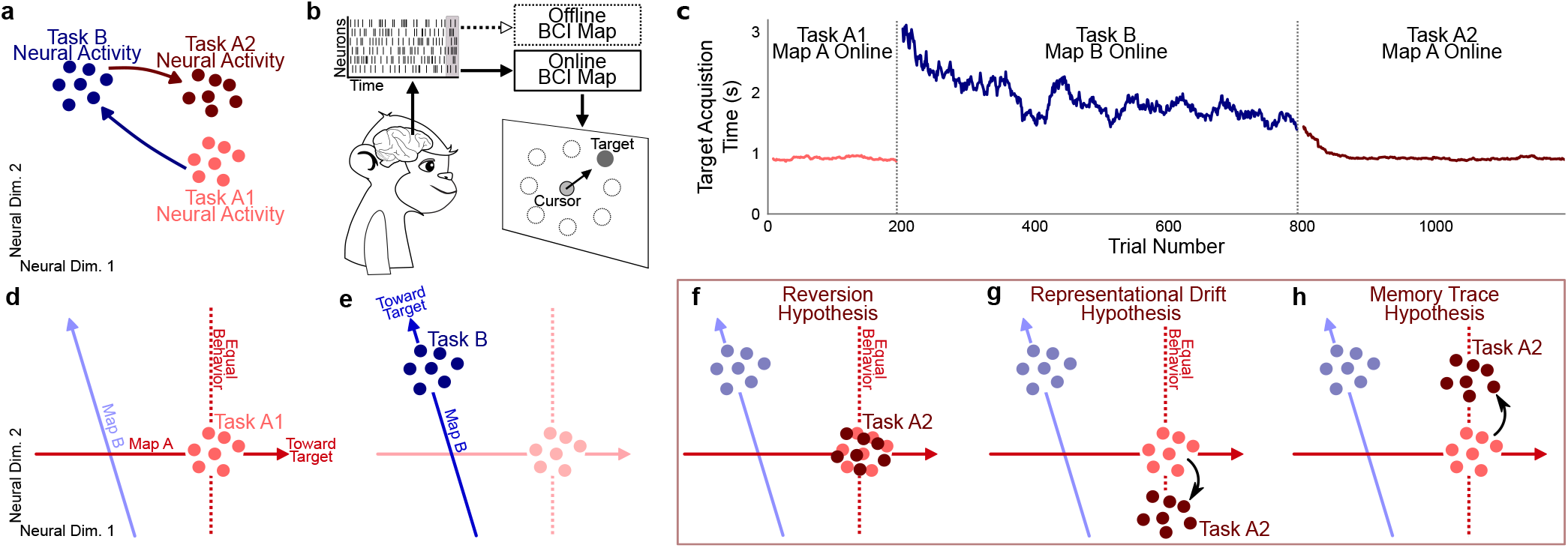
How learning a new task might change the neural activity used for a familiar task. **(a)** Schematic of how neural activity (colored dots) may change when performing different tasks. Performing Task A for the first time (light red; Task A1), then Task B (blue, Task B), then Task A again (dark red; Task A2) may all yield distinct neural activity patterns. **(b)** The activity of ∼90 neural units, recorded using a Blackrock array implanted in the primary motor cortex (M1), were translated into the movement of the cursor through a brain-computer interface (BCI). A BCI directly relates neural activity to behavior (the horizontal and vertical velocities of a cursor on a computer screen) using a map specified by the experimenters. The online BCI map is the BCI map that dictates cursor movements. The same neural activity can also be interpreted with respect to an offline BCI map that did not determine cursor control movement. **(c)** Target acquisition times for an example session (N20160714). The initial period where the monkey used Map A to control the cursor is defined to be Task A1 (acquisition times shown in light red). For Task B, the map was switched to Map B. The monkey had to learn how to control the cursor with this new map through trial and error (dark blue). Acquisition times improved, showing that learning occurred. For Task A2, Map A was reinstated (dark red). For visualization, acquisition times were smoothed with a causal 25-trial moving window and are not shown for the first 8 trials of each task. Success rates were near 100% for all three tasks. **(d-e)** Schematics of how neural activity might look during the three tasks. For illustrative purposes, we show a 2D neural space, which was mapped to a 1D cursor velocity. In the actual experiments, the neural space was ∼90D (one dimension per recorded unit), which was mapped to a 2D cursor velocity. **(d)** During Task A1, neural activity is appropriate for Map A. **(e)** During Task B, neural activity becomes appropriate for Map B. **(f-h)** We explore three possibilities for what neural activity might look like during Task A2. **(f)** Reversion hypothesis: Task A2 neural activity is similar to that used during Task A1. **(g)** Representational Drift Hypothesis: Task A2 neural activity is different from that used Task A1, but not in a manner that consistently retains high performance through Map B. **(h)** Memory Trace Hypothesis: Task A2 neural activity contains a memory trace, whereby neural activity is appropriate for both Map A and Map B.

Our central question is: how does learning Map B affect the neural activity produced while using Map A? To illustrate the possibilities, we depict two dimensions of neural population activity controlling 1D cursor movements (Fig. 1d). During Task A1, the monkey produces neural activity appropriate for Map A, in that the projection of neural activity onto Map A results in high cursor velocities toward the target. During Task B, the monkey learns to produce neural activity that is appropriate for Map B (Fig. 1e). Finally, Map A is reinstated during Task A2, and the monkey’s neural activity needs to once again become appropriate for Map A.

We consider three possibilities for what neural activity might look like after behavior stabilizes during Task A2. One possibility is that, after learning, the population activity patterns produced during Task A2 are similar to those produced during Task A1. We call this the reversion hypothesis (Fig. 1f). Reversion has been observed in various different contexts, such as reaching tasks (Perich and Miller, 2017, Cherian et al., 2013), BCI tasks in visual cortex (Jeon et al., 2022), and in the remapping of hippocampal place fields (Alme et al., 2014). This would indicate that the neural activity we observed in M1 during performance of a task can be unaffected by an intervening learning experience.

A second possibility is that neural activity changes in a manner agnostic to the learning experience. We call this the representational drift hypothesis (Fig. 1g; see Druckmann and Chklovskii (2012), Rule et al. (2019), Mau et al. (2020), Deitch et al. (2021), Schoonover et al. (2021)). Representational drift could occur alongside proficient task performance due to many activity patterns corresponding to the same behavioral output (Kaufman et al., 2014, Hennig et al., 2018). This drift could be attributed to any number of uncontrolled factors, such as arousal (Cowley et al., 2020, Hennig et al., 2021a).

A third possibility is that changes in neural activity are directly related to the learned task. We consider the possibility that neural activity changes to maintain the memory of the learned task (Task B), while simultaneously supporting accurate cursor movement (i.e., action) during Task A2. We call this the memory trace hypothesis (Fig. 1h). Neural activity changing in this manner could help facilitate the formation of new memories without leading to interference with subsequent behavior. While prior work has observed changes in neural activity as a result of an intervening learning experience and speculated that these changes reflect a memory trace (Li et al., 2001, Arce et al., 2010), with a BCI we know the causal relationship between neural activity and behavior and thus are now able to disambiguate between the representational drift and memory trace hypotheses.

We commence our analyses by considering the reversion hypothesis. If the reversion hypothesis were true, we would expect the tuning of individual neural units to remain the same between Tasks A1 and A2. To test this, we fit cosine tuning curves in each of these task periods and measured the change in preferred direction between them. We found many neurons exhibited substantial tuning changes (Fig. 2a). Overall, these tuning curve changes confirm that neural activity produced during Task A2 is distinct from that of Task A1 at the single unit level (Fig. 2b). A lack of support of the reversion hypothesis is also evident when we consider the population of neurons together (Fig. 2c). We observed that, for many targets, neural activity during Tasks A1 and A2 occupied different regions within the neural population space (Fig. 2d), in contradiction to the schematic in Fig. 1f. Thus, our data are not consistent with the reversion hypothesis.

**Figure 2.**
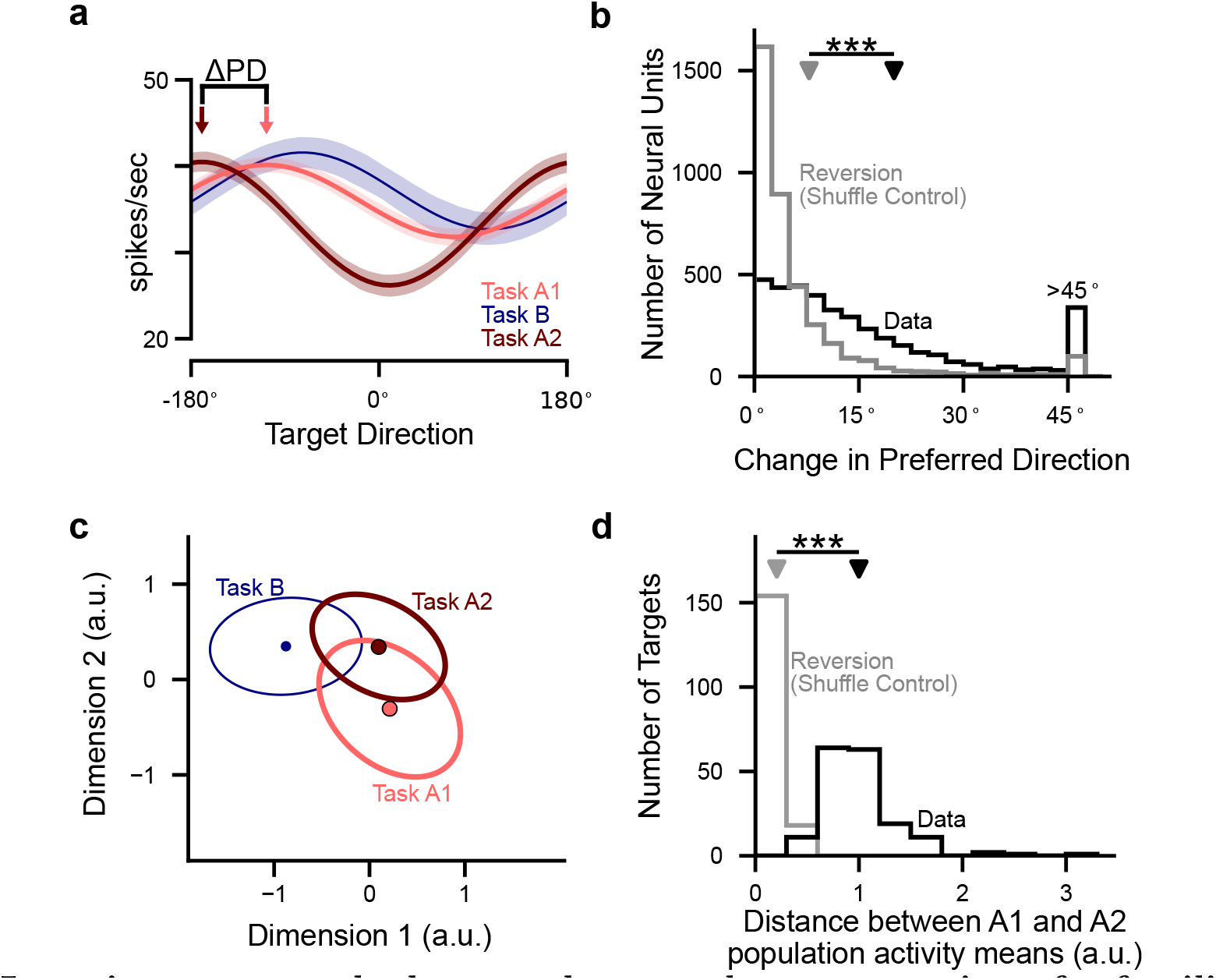
Learning a new task changes the neural representation of a familiar task. **(a)** Tuning curves relating cursor-to-target direction to the firing rate for an example neural unit. A cosine tuning curve was fit separately for each of the three task periods. This unit (unit 37 from session L20131205) changes its tuning (measured by a change in preferred direction, Δ PD) between Tasks A1 and Tasks A2. Shading indicates a 95% confidence interval. **(b)** Many units show a change in tuning between Tasks A1 and A2 (*P <* 10^−10^, two-sided paired Wilcoxon signedrank test, n=3461 neural units). Black shows the absolute change in preferred direction for units across all sessions. Grey indicates the prediction of the reversion hypothesis (that is, no change in PD other than that due to sampling error). This was estimated using a shuffle control in which labels for Task A1 and A2 were randomly permuted across trials (see Methods). **(c)** A view of the population neural activity for one example target (J20120601; target 270°) across all three task periods. We applied linear discriminant analysis (LDA) to find the plane which best separates the neural activity from the three task periods. Activity is projected onto that plane, with mean and covariances across timesteps shown. **(d)** Population activity is different between Task A1 and Task A2 (*P <* 10^−10^, two-sided paired Wilcoxon signed-rank test, n=172 targets). Black shows the distance between the Task A1 and Task A2 means in the 10D population activity space. Grey indicates the prediction of the reversion hypothesis, obtained using a shuffle control (see Methods).

Although we can rule out the reversion hypothesis, our analyses to this point does not distinguish the memory trace hypothesis from the representational drift hypothesis. To do so, we must evaluate how the observed changes in neural activity relate to the previously-learned behavior. Our BCI approach makes this possible because we can quantify whether the neural activity is suitable for a BCI map that is not currently being used by the monkey. To illustrate this process, we compare neural activity from a single trial during each of Task A1 and Task A2 corresponding to the same target (Fig. 3a top). For each population activity pattern, we can evaluate its “progress” through Map A as the extent to which it moves the cursor toward the target (see Methods). During both Tasks A1 and A2, Map A determines cursor velocity, and the monkeys showed proficient control of the cursor during both tasks (Fig. 3b; Extended Data Fig. 1).

**Figure 3.**
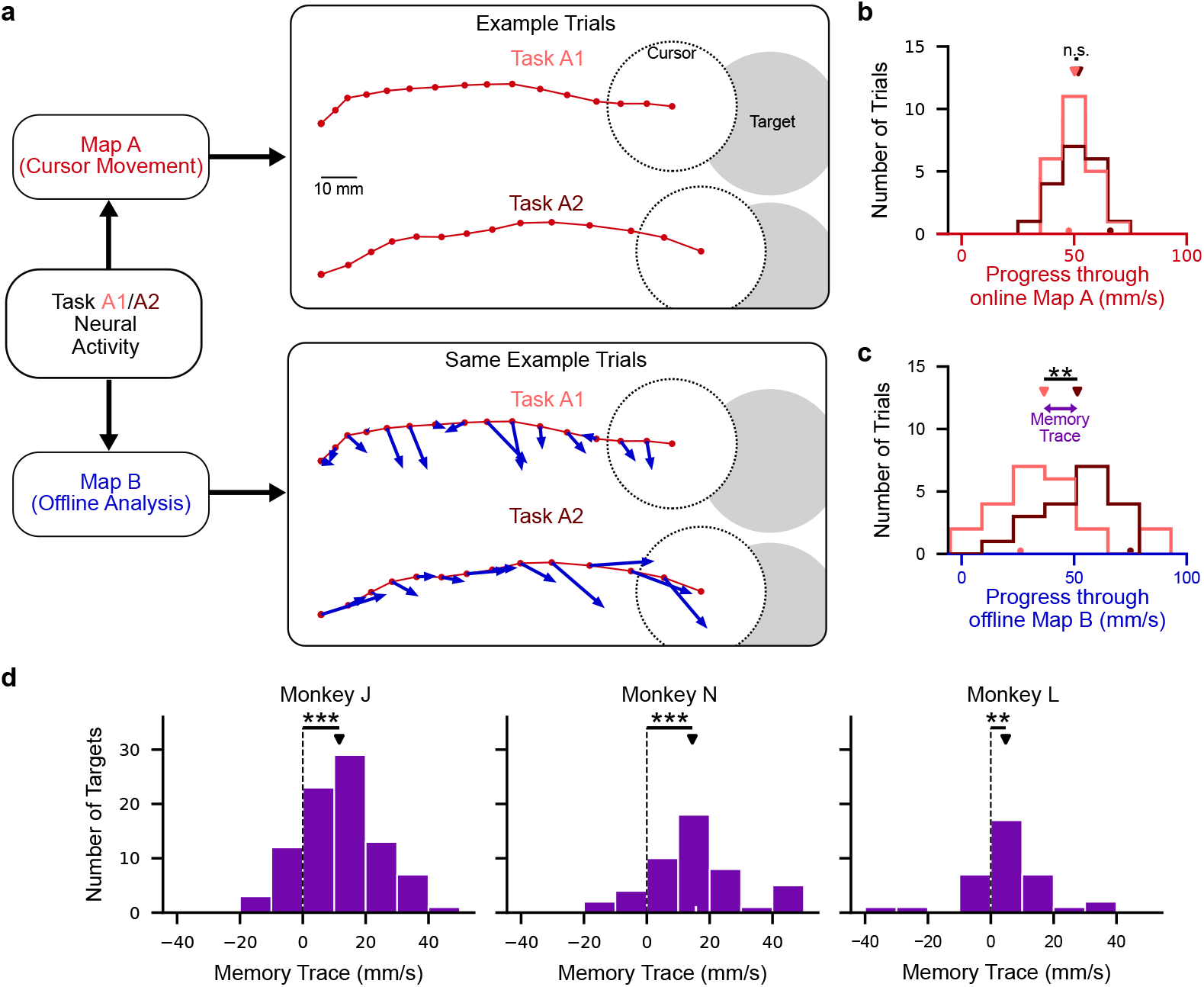
Learning leaves a memory trace. **(a)** During Task A1 and Task A2, neural activity drives the cursor through Map A (red trajectories, with dots denoting cursor positions at each timestep). The same neural activity can also be projected through Map B in an offline analysis (blue arrows). During Task A2, the Map B velocities more directed toward the target than during Task A1. Both trials come from the 225° target from session N20160329. For visualization purposes, the data are rotated and the velocities are scaled. **(b)** Task performance is similar between Task A1 and Task A2 (see Extended Data Fig. 1). For this target, there is no significant difference in progress (i.e., the component of velocity that points toward the target), through Map A (*P* = 0.80, two-sided unpaired Wilcoxon rank-sum test). Dots on the horizontal axis denote the average progress for the trials shown in **(a)**. Triangles above the histograms denote the mean of each distribution. **(c)** Velocities through the offline Map B. The difference in average progress defines the memory trace for that target. For this target, there is significantly higher progress through Map B during Task A2, relative to Task A1 (*P* = 0.0077, two-sided unpaired Wilcoxon rank-sum test) yielding a memory trace of 14.49 mm/s. Same conventions as in **(b). (d)** All three monkeys showed a memory trace for well-learned targets (Monkey J, *P <* 10 × 10^−10^, two-sided paired Wilcoxon signed-rank test, n=88 targets; Monkey N, *P* = 1.14 × 10^−7^, n=48 targets; Monkey L *P* = 0.0020, n=36 targets). For a small fraction of targets, the measured memory trace is negative. This arises when progress through Map B is worse during Task A2 than Task A1. When also including unlearned targets, a memory trace is still evident for Monkey’s J and N, but not Monkey L (Monkey J, *P* = 1.57 × 10^−4^, two-sided paired Wilcoxon signed-rank test, n=176; Monkey N, *P* = 1.99 × 10^−6^, n=96; Monkey L *P* = 0.61, n=72; see Extended Data Fig. 3). Monkey L showed a smaller memory trace than Monkeys J and N, likely due to less learning occurring (Extended Data Fig. 4). Triangles denote the average memory trace for each monkey. The white tick mark on the horizontal axis of the middle histogram denotes the example target illustrated in **(b)** and **(c)**.

Since we are using a BCI, progress can also be calculated for Map B, even when the animal is using Map A to control the cursor. Progress under Map B measures the extent to which a given neural activity pattern *would have moved* the cursor toward the target, had Map B been instantiated. During Task A1, the monkeys exhibited low progress through Map B, as the velocities through Map B are small and haphazardly oriented relative to the target (Fig. 3a, bottom, Task A1). This is expected because the monkey had not yet experienced Map B, and Map B was selected to be difficult to control using Map A’s neural activity (Sadtler et al., 2014). In contrast, during Task A2 the velocities through Map B are larger and more directed toward the target than they were during Task A1 (Fig. 3a, bottom, Task A2), that is, they show higher progress (Fig. 3c). This occurs despite the fact that Map B has no influence on behavior during Task A2 and thus the monkeys have no external incentive while performing Task A2 to maintain high progress through Map B. We define the memory trace as the average increase in the progress toward a given target when projecting the neural activity patterns through Map B during Task A2, relative to Task A1. Across all three monkeys, we found that average progress through Map B was larger during Task A2 than Task A1 (Fig. 3d). This finding supports the memory trace hypothesis (Fig. 1h), but not the representational drift hypothesis, which does not predict this organization.

We next assessed the robustness of the memory trace with two tests. First, we showed the memory trace was still present when using a different performance metric, namely, angular error (Extended Data Fig. 2). Second, we quantified the consistency of the effect by showing that the majority of targets from each session exhibited a memory trace, and that the average of the memory traces per session is positive (Extended Data Fig. 3).

We next considered whether the memory trace possesses three desirable properties of useful memories. The first property is that a memory should *persist*, meaning that it is present in neural activity without dissipating as time passes. To test this, we examined the later trials of the sessions with the longest Task A2 blocks (Fig. 4a). Specifically, we split sessions into two groups. The first group contained the sessions with at least 300 Task A2 trials, while the second group contained sessions with fewer than 300 Task A2 trials (see Methods). For the group with the longer Task A2 period, we excluded the first 200 trials from analysis in order to quantify the memory trace after extended usage of Map A. We found that the memory trace at the end of these longer sessions was not detectably different from the memory trace of the sessions with fewer Task A2 trials (Fig. 4b).

**Figure 4.**
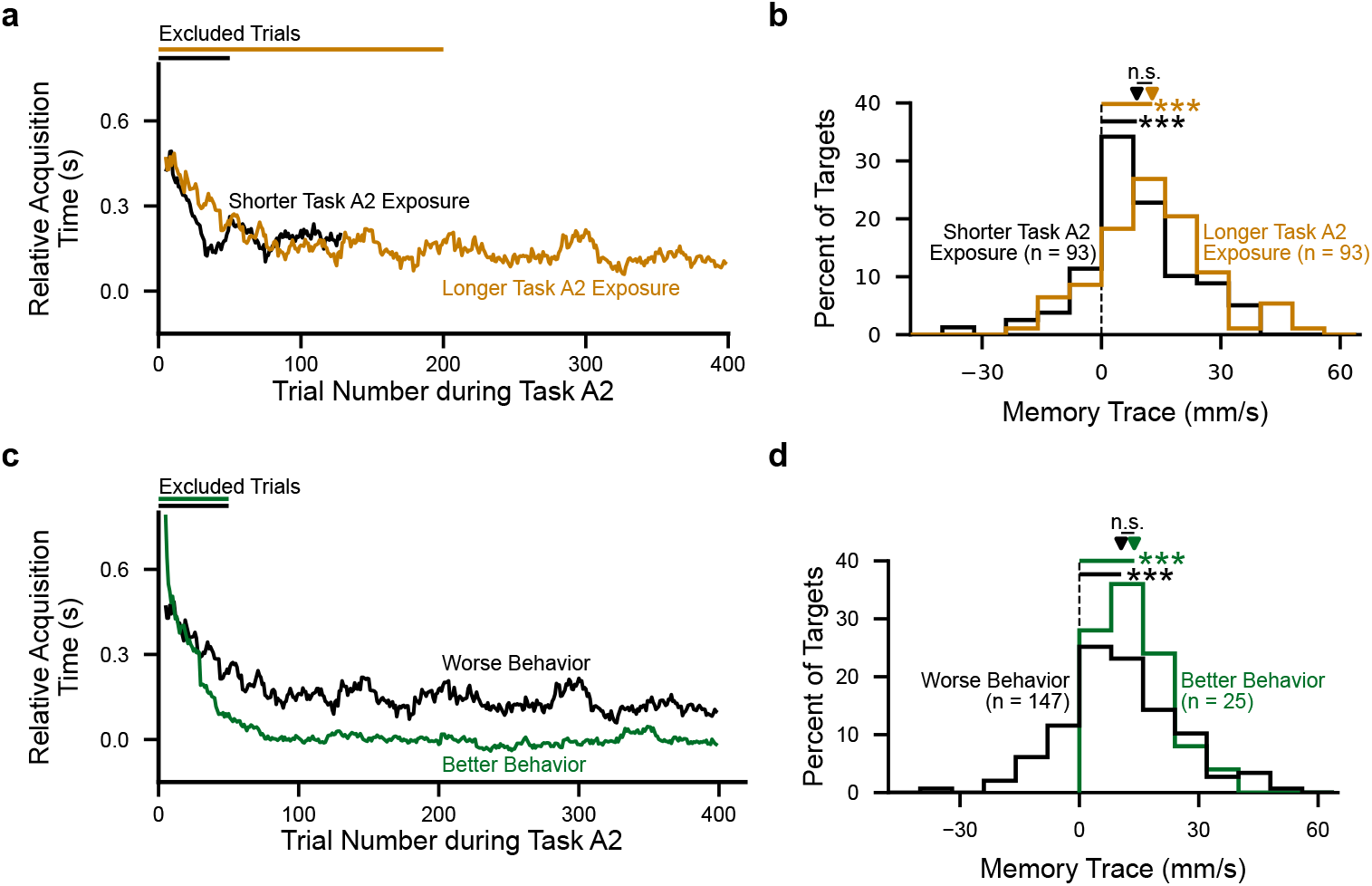
The memory trace persists over time and coexists alongside proficient task performance. **(a)** To examine the influence of longer Task A2 exposure on the memory trace, we split sessions into two groups. The first group contains sessions with fewer Task A2 trials (behavioral performance for an example session shown in black), while the second group contains sessions with more Task A2 trials (example session in gold). For the group of shorter sessions, we excluded the first 50 Task A2 trials (black bar). For the longer group, we excluded the first 200 trials (gold bar). Acquisition times are plotted relative to the Task A1 period, where zero represents the an acquisition time equal to the average acquisition time for that target during Task A1. **(b)** The size of the memory trace was not different for shorter versus longer Task A2 exposure (*P* = 0.11, two-sided unpaired Wilcoxon rank-sum test). The memory trace was still evident for both the longer sessions (gold; *P <* 10^−10^, two-sided paired Wilcoxon sign-rank test), and the shorter sessions (black; *P* = 2.02 × 10^−8^). **(c)** Behavioral performance during Task A2 from two example sessions, one session with better behavior (faster acquisition time; green) and the other with worse behavior (slower acquisition times; black). Note that the session with worse behavior is the same session as that shown in **(a)** for the longer Task A2 exposure. **(d)** To evaluate the influence that behavioral performance during Task A2 has on the size of the memory trace, we split targets into two groups. The first group contained targets where the mean target acquisition time during Task A2 was less than the mean target acquisition time during Task A1 (“better behavior”; green; see Extended Data Fig. 1), The second group contained targets where the mean target acquisition time during Task A2 was greater than during Task A1 (“worse behavior”; black). The size of the memory trace was not different between the worse behavior and better behavior groups (*P* = 0.12, two-sided unpaired Wilcoxon sign-rank test). Moreover, the memory trace was still evident in the group of targets with better behavior (*P* = 5.96 × 10^−8^, two-sided paired Wilcoxon rank-sum test) and worse behavior (*P <* 10^−10^).

The second desirable property of a memory is that it should *coexist* alongside proficient performance of other tasks. To address this, we examined whether the size of the memory trace was contingent on how proficient the behavior was during Task A2 (Fig. 4c). If the instances with worse behavioral performance during Task A2 had the largest memory trace, it could suggest that the memory trace arises due to a trade-off between performance through the two BCI maps. Alternatively, if the memory trace were present even when behavioral performance during Task A2 returned to the levels seen during Task A1, it would suggest that the memory trace can coexist without hindering the monkey’s ability to perform the familiar task. We found that the memory trace for the targets with the best behavioral performance during Task A2 showed an average memory trace whose strength was not significantly different from the average memory trace of targets with worse behavioral performance during Task A2 (Fig. 4d). These results indicate the memory trace coexists alongside proficient behavioral performance of the familiar task, and does not represent a compromise between the two learned behaviors.

The final property is that more learning should lead to more memory. We found that to be the case, as the size of the memory trace was positively correlated with the amount of learning during Task B (Extended Data Fig. 4). As Monkey L showed less learning than the other two monkeys, this could explain why its memory trace tended to be smaller (Fig. 3d, Monkey L).

How can a memory trace coexist without degrading behavioral performance during Task A2? To understand this, we considered how the changes in neural activity induced by learning Map B relate to Map A. Because we map the activity of ∼90 neural units to two-dimensional BCI cursor movements (see Methods), not all changes in neural activity affect cursor movement. We refer to changes in neural activity that affect cursor movement as “output-potent” with respect to that map, and changes that do not as “output-null”(Kaufman et al., 2014). Because Map A and Map B do not share the same output-potent space, it is possible to have neural changes that affect cursor movement through one map without impacting cursor movements through the other.

We examined whether the memory trace of Map B (Fig. 5a) resides in the outputpotent or output-null space of Map A (Fig. 5b), by decomposing it into its output-potent and output-null components (Fig. 5c). We found that the memory trace resides predominantly in the output-null space of Map A (Fig. 5d), rather than in the output-potent space of Map A (Fig. 5e). This means the memory trace is primarily “stored” in dimensions that do not influence task performance (Extended Data Fig. 5). Since neural activity in dimensions output-null to Map A do not influence cursor velocities during Task A2, this explains how the memory trace can co-exist with proficient behavioral performance.

**Figure 5.**
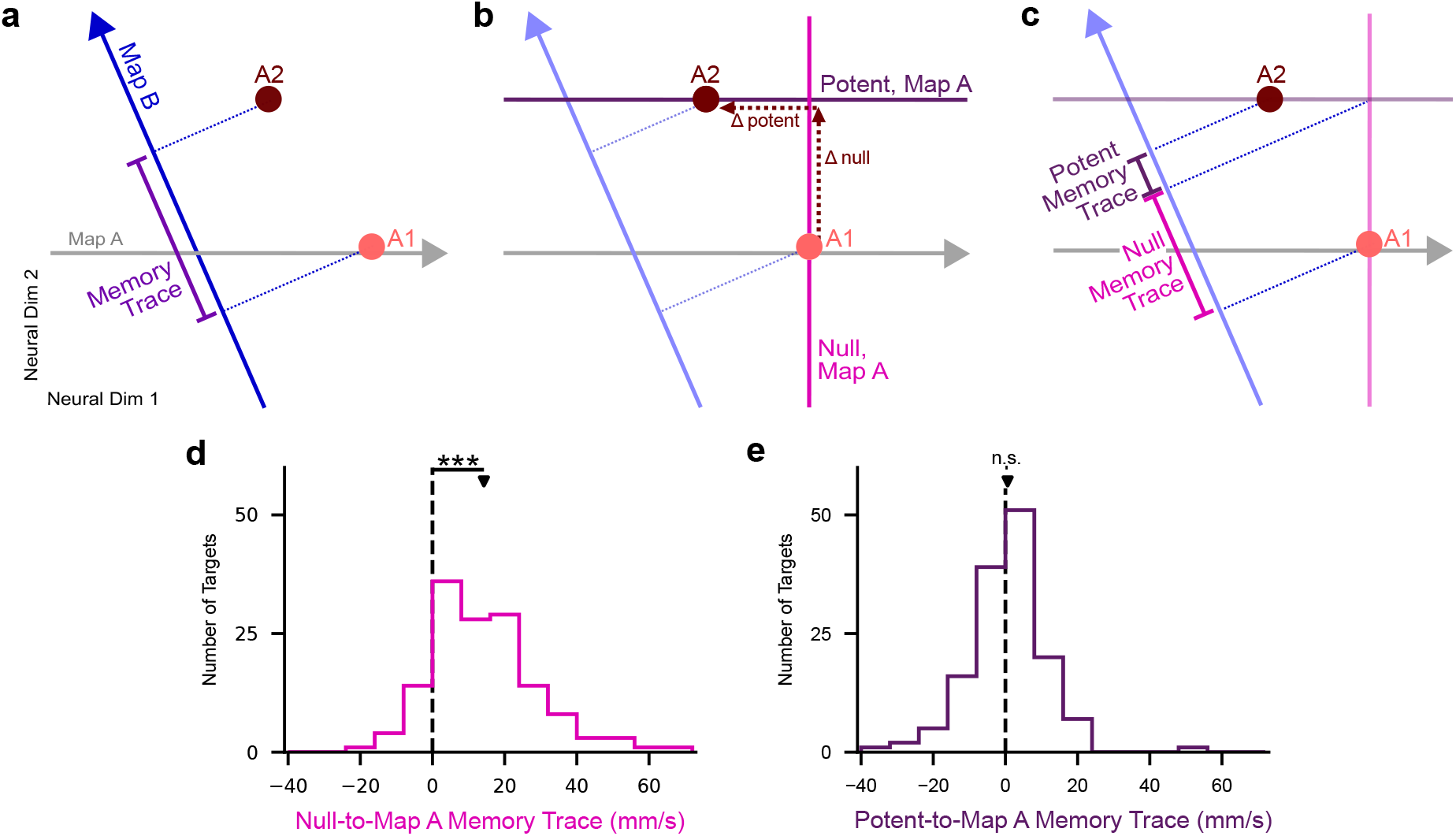
The memory trace is predominantly in the null space of Map A. **(a)** Memory trace depicted in same space as Fig. 1d-h. Between Task A1 (light red dot) and Task A2 (dark red dot), neural activity changes. During these time periods, the cursor is controlled using Map A (grey arrow). Task A2 activity is further along Map B (blue arrow) than Task A1 activity, indicating higher progress. The memory trace is defined as difference in the projection onto Map B. **(b)** The change in neural activity from **(a)** can be decomposed into a component that is output-potent to Map A (Δ potent) and a component that is output-null to Map A (Δ null). **(c)** Having decomposed the change in neural activity into output-potent and output-null components, we can correspondingly decompose the memory trace into output-potent and output-null components. **(d)** Of the targets with a positive memory trace (142 out of 172 targets), the memory trace consistently resides in dimensions null to Map A (*P <* 10^−10^, two-sided paired Wilcoxon signed-rank test, n=142 targets across all monkeys). **(e)** The contributions from the potent space are not significantly different from zero (*P* = 0.31, two-sided paired Wilcoxon signedrank test, n=142 targets across all monkeys), meaning there is no memory trace on average in the output-potent component of neural activity.

Last, we considered, how does the monkey arrive at the Task A2 solution? There are two possibilities. The first possibility is that there is a partial “unwinding” of the learning that occurred during Task B. This would suggest that the solution used during Task A2 is not novel, and was employed sometime during the learning experience. If this were true, we would expect that the path neural activity takes from the end of Task B to the end of Task A2 (i.e., “the path of washout”, dark red arrow in Fig. 6a) would retrace the path that neural activity takes from the end of Task A1 to the end of Task B (i.e., “the path of learning”, blue arrow in Fig. 6a). The other possibility is that the path of washout is distinct from the path of learning (Fig. 6b). This would imply that the solution the monkey uses during Task A2 is novel, suggesting that the relearning of Map A is distinct from simply “forgetting Map B”. To differentiate between these possibilities, we calculated the angle formed between the path of learning and the path of washout (see Methods). We found that the path of washout is distinct from the path of learning (Fig. 6c).

**Figure 6.**
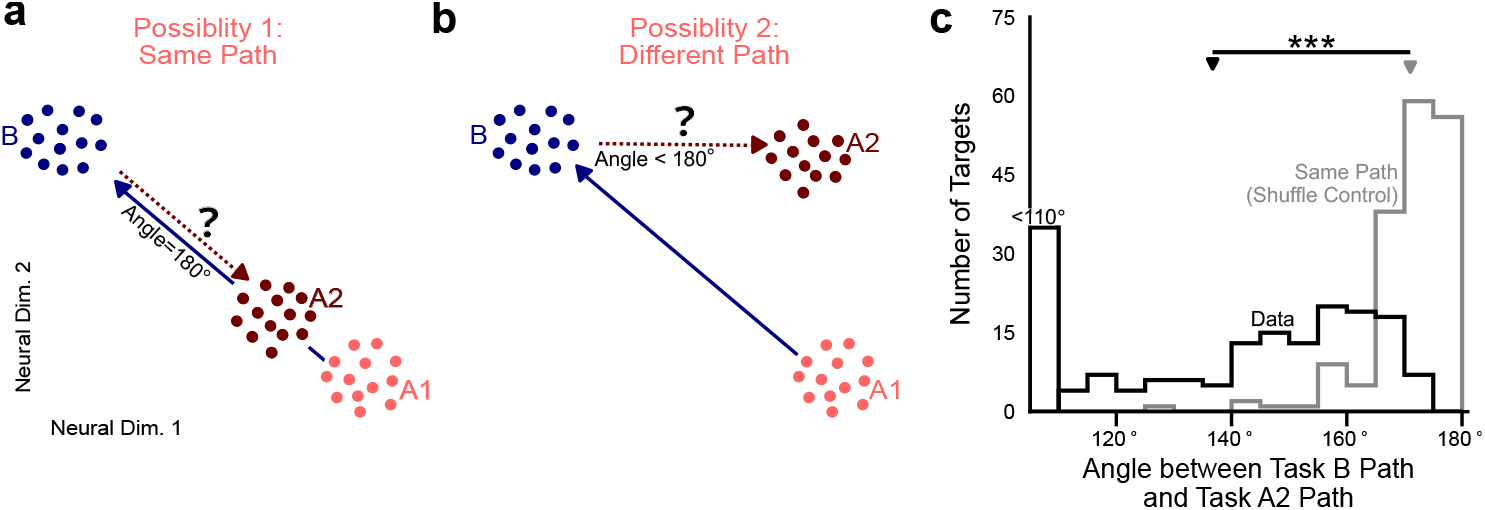
The path of washout does not retrace the path of learning. We consider two possibilities for how the memory trace arises during Task A2. **(a)** The first possibility is that the path of washout (i.e., the path neural activity takes from the end of Task B to Task A2) retraces the path of learning (i.e., the path neural activity takes from Task A1 to the end of Task B). This would mean that washout is simply an “unwinding” of the learning experience. (b) The second possibility is that these two paths are distinct, implying that the washout is not simply “unlearning”. **(c)** To distinguish between these two possibilities, we measured the angle between these two paths. The angle between these paths (black histogram) was smaller than the angles that would be obtained under possibility 1 (grey histogram; see Methods; *P <* 10^−10^, twosided paired Wilcoxon signed-rank test, n=172 targets across monkeys). This implies the paths of learning and washout are distinct (possibility 2).

## Discussion

We studied how the brain can retain a memory of a newly learned task without compromising the performance of familiar tasks. A BCI enables us to make new progress on this longstanding question. This is because using a BCI allows us assess the extent to which the same neural population activity patterns were suitable for multiple different maps (that is, relationships between neural activity and cursor movements), including a map not actively being used. We found that, after a learning experience, neural activity remained appropriate for the learned map even when the animal was using a different (familiar) map. The memory of the learned map was primarily in dimensions in neural space which were output-null to the familiar map. In this way, neural activity simultaneously supported action through the familiar map and still maintained a memory of the recently learned map.

It could have been that motor memories were stored in a manner that is not detectable when another action is being performed (Herzfeld et al., 2014, Jeon et al., 2022), nor be present in the same neural activity when it is driving behavior. For example, in our experiments, the motor memory could have been stored (perhaps outside of M1) such that the memory is only detectable in M1 the appropriate behavior is being performed. Instead, we found that memories can be stored in a manner that makes them evident in M1 even during the execution of other actions.

Motor memory consolidation is the process by which memories become more robust to interference (Krakauer et al., 2005). This process takes at least several hours (Shadmehr and Holcomb, 1997), and may require M1 (Muellbacher et al., 2002, Kawai et al., 2015, Rubin et al., 2022). How might the brain bridge from the short-timescale retention of a memory trace that we studied here to the longer-timescale consolidation of a motor memory (Shadmehr and Holcomb, 1997, Gulati et al., 2017)? Our results focused on the short-term inception of a motor memory, within an hour or so of the learned experience. Three possibilities would be consistent with our results. First, a long-term consolidated memory might resemble the memory trace we observed here. Second, it might be that the memory trace we observed is only a short-term phenomenon in M1, dissipating after consolidation. Thus, the memory trace evident in M1 could constitute a short-term storage for the memory before it is offloaded to another brain area during consolidation. Finally, it could be that with further practice with both maps over many days, the neural activity changes to provide even better performance through both BCI maps. In this way, the monkey could effortlessly switch between the two tasks without a drop in performance using the same population activity patterns. That is, with further practice the memory trace could evolve (Nader and Hardt, 2009, Gershman et al., 2017) to lead to even greater coexistence between the two behaviors (Ajemian et al., 2013, Gallego et al., 2018).

What is the utility of maintaining a memory trace in neural population activity? A memory trace could enable proficient performance to be reached more quickly upon re-exposure to the learned task. This phenomenon, known as savings, has been frequently observed in motor learning behavior and is often taken as evidence of memory formation (Krakauer et al., 2005, Herzfeld et al., 2014). Our results propose a neural population mechanism for savings. Namely, if Map B were to be re-introduced following performance of Task A2, neural activity would already be situated in population activity space in a manner that would yield better initial performance while using Map B than before learning. While this mechanism can lead to savings due to starting from a better position, our results do not speak to whether there would also be an increased rate of relearning, i.e., a greater reduction in error per trial after the first trial.

The memory trace we found in M1 represents one scheme whereby the brain can store multiple memories without interference. We found that the firing of many neurons contribute to the memory trace. This coding scheme marks an interesting contrast to how some memories are formed in the hippocampus, where a sparse subset of neurons encode the memory (Josselyn and Tonegawa, 2020). We observed that the memory trace was primarily due to changes in neural activity orthogonal (i.e., output-null) to the familiar task. Notably, the utilization of different subsets of neurons to encode memories is a special case of orthogonal representations in population activity space (Alme et al., 2014). These lines of evidence together indicate that the brain needs to incorporate new memories into subspaces orthogonal to existing memories in order to avoid interference (Ajemian et al., 2013, Tang et al., 2020, Gava et al., 2021, Libby and Buschman, 2021, Nieh et al., 2021, Xie et al., 2022). Avoiding interference may be harder in the spinal cord, where there are fewer neurons than in cortex or the hippocampus. As fewer neurons likely leads to a more constrained encoding space, a “negotiated equilibrium” between multiple learned behaviors may be required (Wolpaw, 2018).

By demonstrating the presence of a memory trace, we ruled out the possibility that changes in neural activity between Task A1 and Task A2 were due solely to representational drift, a change in neural activity manner agnostic to the learning experience. However, representational drift has been observed throughout the brain (Druckmann and Chklovskii, 2012, Rule et al., 2019, Mau et al., 2020, Deitch et al., 2021, Schoonover et al., 2021), and could be occurring alongside the memory trace that we observe. Representational drift differs from the formation of a memory trace in that the changes in neural activity due to representational drift do not directly serve the purpose of memory, but instead are driven by other factors, not under experiment control.

Sun et al. (2022) also observed systematic changes in neural activity related to the learning experience. In their study, learning an arm-reaching task in a curl force field induced a uniform shift in preparatory neural activity, which persisted after the force field was removed. The authors conjecture that this shift indexes motor memories (Sheahan et al., 2016). It remains to be seen whether uniform shifts during preparatory activity lead to the reorganization of activity in M1 that constitutes a memory trace, or if these findings support two separate processes.

Human and animal learners distinguish themselves from current artificial learning systems in that they can learn to perform a large number of different behaviors. It is a notoriously challenging problem for artificial agents to learn new tasks without overwriting the ability to perform previously learned tasks, an effect termed “catastrophic forgetting” (Masse et al., 2018, Parisi et al., 2019, Yang et al., 2019). Our findings suggest that artificial learning systems could overcome catastrophic forgetting by implementing some of the same learning principles employed by biological learning systems (Duncker et al., 2020, Hennig et al., 2021b). A sufficiently high dimensional activity space may be important not only in the brain, but also for artificial agents, for learning multiple tasks without interference.

## Acknowledgments

We thank Caleb Kemere for helpful discussions. This work was supported by the National Science Foundation Graduate Research Fellowship DGE1745016 and DGE2140739 (to D.M.L.), the Richard King Mellon Presidential Fellowship (to J.A.H.), the Carnegie Prize Fellowship in Mind and Brain Sciences (to J.A.H.), NIH R01 HD071686 (to A.P.B., B.M.Y. and S.M.C.), NSF NCS BCS1533672 (to S.M.C., B.M.Y. and A.P.B.), NSF NCS DRL 2124066 and 2123911 (to B.M.Y., S.M.C. and A.P.B.), NSF CA-REER award IOS1553252 (to S.M.C.), NIH CRCNS R01 NS105318 (to B.M.Y. and A.P.B.), NSF NCS BCS1734916 (to B.M.Y.), NIH CRCNS R01 MH118929 (to B.M.Y.), NIH R01 EB026953 (to B.M.Y.), Simons Foundation 543065 (to B.M.Y.), NIH R01 NS120579 (to A.P.B.) and NIH R01 HD090125 (to A.P.B.).

## Author Contributions

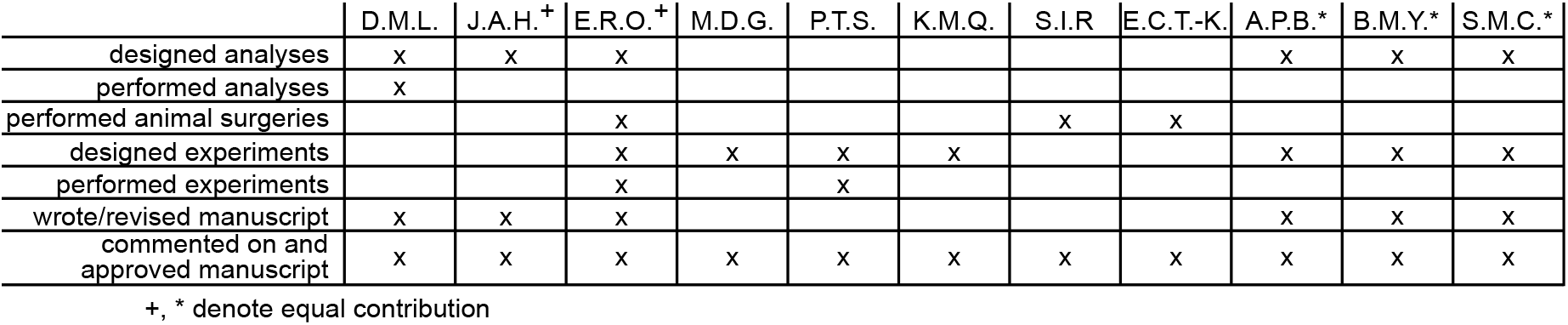

## Methods

### Experimental procedures

Experimental methods are detailed in our previous work (Sadtler et al., 2014, Golub et al., 2018). Briefly, we recorded neural activity from three male Rhesus macaques (Maccaca mulatta, ages 7, 7 and 8 for monkeys J, N and L, respectively) using 96 electrode arrays (Blackrock Microsystems) implanted in the proximal arm region of the primary motor cortex. All animal care and handling procedures conformed to the NIH Guidelines for the Care And Use of Laboratory Animals and were approved by the University of Pittsburgh’s Institutional Animal Care and Use Committee.

The monkeys performed an eight-target center-out BCI task. In the BCI, a monkey guided a computer cursor by modulating its neural activity. The recorded neural activity was translated into movements of the computer cursor according to a BCI map (see *Translating neural activity to cursor movement*). Each session was split into three task periods, “Task A1”, “Task B”, and “Task A2”. The three task periods followed the same experimental paradigm, differing only in the BCI map. During Task A1 the monkey used Map A, which was selected to be intuitive for the monkey to use from the outset. The monkey controlled the cursor during Task A1 for 318.8 ± 95.4 (mean ± s.d.) trials. Uncued to the monkey, we then switched to Map B for the second period of the experiment (Task B). The monkey had never seen before Map B and was selected in order to initially be difficult for the monkey to use to control the cursor. The monkey was given 696.7 ± 219.4 (mean ± s.d.) trials to learn to control the cursor with Map B. Finally, again uncued, Map A was reinstated (Task A2). The Task A2 period lasted the remainder of the experiment for 318.2 ± 153.9 (mean ± s.d.) trials.

#### Trial flow

At the start of each trial, the cursor appeared at the center of the monkey’s workspace. Target locations were selected pseudo-randomly from a set of eight uniformly spaced locations around a circle (radius, Monkey J: 150 mm; Monkeys L and N: 125 mm). The target appeared on the screen at the beginning of the trial. For the first 300 ms, the cursor’s velocity was fixed at zero. After this, the velocity of the cursor was controlled by the monkey through the BCI map corresponding to the task period of the experiment. If the monkey was able to acquire the target within 7.5s after the start of the trial, a water reward was dispersed. If the monkey failed to acquire the target within the allotted time, there was a 1.5s timeout prior to the start of the next trial.

#### Identifying latent dimensions of neural activity

Experiments began with a calibration period in order to define Map A. Monkey J’s calibration employed either passive cursor observation or closed-loop BCI control using the previous day’s BCI map. For monkeys L and N, we used a calibration procedure that gradually stepped from passive observation to closed-loop control. We then applied factor analysis (see below) to identify the 10D linear subspace (the “intrinsic manifold”) that captured the dimensions of greatest shared variance in the neural population. Ten dimensions was selected using cross-validation, as described in prior work (Sadtler et al., 2014). Spike counts (i.e. threshold crossings) were taken in nonoverlapping 45 ms time windows. We denote the spike counts at timestep t as **u**_*t*_ ∈ ℝ^*q×*1^, where *q* is the number of neural units. Factor analysis describes this high-dimensional population activity, **u**_*t*_, in terms of a low-dimensional set of factors, **z**_*t*_ ∈ ℝ^10×1^. Latent factors, **z**_*t*_, are distributed as:

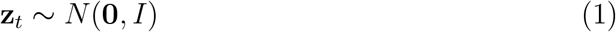

where *I* is the identity matrix. Spike counts, **u**_*t*_, are related to the factors by:

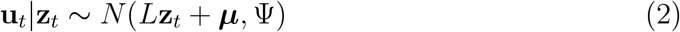

where parameters *L* ∈ ℝ^*q×*10^ (termed the loading matrix), ***µ*** ∈ ℝ^*q×*1^, and Ψ ∈ ℝ^*q×q*^ (a diagonal matrix of variances independent to each neuron) are estimated using the expectation-maximization algorithm. The latent factor activity, **z**_*t*_, at timestep *t* is estimated as the posterior expectation given the spike counts as:

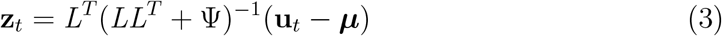

For all analyses, we orthonormalized **z**_*t*_ so that it had units of spike counts per timestep to facilitate the interpretability of the factor activity. As the majority of the shared variance of the neural population is captured in these latent dimensions, and neural activity cannot be readily produced outside this low-dimensional subspace during short-term learning (Sadtler et al., 2014, Oby et al., 2019), we focus our analyses on this factor activity, referred to as “population activity patterns” throughout.

#### Translating neural activity to cursor movement

At each 45 ms timestep *t*, neural activity drove the computer cursor according to the BCI map for that task period. Specifically, the cursor velocity was determined using a Kalman filter:

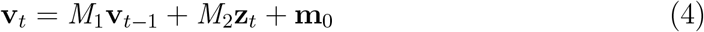

The parameters *M*_1_ ∈ ℝ^2×2^, *M*_2_ ∈ ℝ^2×10^ and **m**_0_ ∈ ℝ^2×1^ are determined during the calibration period (see Sadtler et al. (2014) for details), and **v**_*t*_ ∈ ℝ^2×1^ comprises the horizontal and vertical cursor velocities. The two BCI maps differ only in the *M*_2_ term. For Map A, 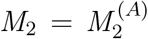, which is found during the calibration session. For Map B, 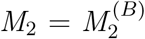 was a permutation applied to the columns of 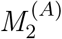, equivalent to permuting the elements of **z**_*t*_ before applying equation 4. This means that Map B remained within the intrinsic manifold (a “within-manifold perturbation”). Thus Map B changed the relationship between the factor activity and cursor velocity.

### Data Analysis

The data analyzed in this study was part of a larger study that included both within-manifold perturbations (WMPs) and outside-manifold perturbations (OMPs) (Sadtler et al., 2014). As we have previously found that WMPs show stronger learning than OMPs, we only considered sessions that used WMPs.

Data from the Task A1 and Task B periods of these WMP sessions were analyzed in prior work (Golub et al., 2018, Hennig et al., 2018, 2021a). Here we focused on neural activity recorded during Task A2, which has not been previously studied. To ensure an adequate amount of Task A2 data to analyze per session, we only considered sessions that included at least 100 Task A2 trials. This yielded a total of 43 sessions (Monkey J, 22 sessions, 362.6 ± 170.2 Task A2 trials; Monkey N, 12 sessions, 333.3 ± 107.3 Task A2 trials; Monkey L, 9 sessions, 171.0 ± 49.7 Task A2 trials; all values mean +/-s.d.).

#### Selecting experiments and trials for analysis

Some targets showed more learning than others. As the focus of this work is on the memory of a learned task, we analyzed targets that showed the most learning. We defined *learning* as how well the monkey performed with Map B after learning, relative to how well it would have performed with Map B if it continued producing the same neural activity as it did during Task A1 (i.e., if there was no learning). Thus, we defined learning as the difference in the average Map B progress (see *Quantifying the memory trace* for how progress is computed) of the last 10 trials to a given target during Task B compared to average Map B progress to that same target during Task A1. For each monkey, the 50% of targets with the most learning were designated as “well-learned.” Well-learned targets had an average amount of learning of 26.61 ± 13.07 mm/s, compared to 0.69 ± 8.79 mm/s for the other targets. Fig. 2a, Fig. 2b, Fig. 4a, Fig. 4c, Extended Data Fig. 3 and Extended Data Fig. 4 include all targets. All other analyses focus on the well-learned targets.

As our central question focuses on neural activity during proficient Task A2 performance, we restricted analyses of Task A2 to after behavior had stabilized. To do this, we excluded the first 50 trials of Task A2 from each session (see Fig. 4). Unless stated otherwise, the remaining Task A2 trials are referred to as Task A2 throughout the manuscript. Additionally, we only analyzed successful trials, as it is otherwise difficult to determine whether the monkey was engaged in the task.

On each trial, we discarded the first 90 ms (2 timesteps during the freeze period) as the activity in M1 would not yet reflect the target due to sensory processing delays (Golub et al., 2015). Additionally, because we report trial-averaged and targetaveraged quantities, we wanted to ensure neural activity came from instances in which the monkey needed to push the cursor in the same direction. Thus, we only analyzed timesteps in which the angle between the cursor and the target was no greater than 22.5° away from the target direction for that trial. Performing our analyses without this exclusion criterion did not change our results qualitatively.

Even after learning to use Map B, the monkeys generally exhibited lower performance with Map B than Map A (see Fig. 1c). Thus, Task B trials tended to be longer than the Task A1 and A2 trials. To compare the Task B trials to the Task A1 and A2 trials, we only utilized the first 25 timesteps from each trial. This number was selected because it is approximately equal to the average Task A1 acquisition time across all monkeys.

#### Testing the reversion hypothesis

To measure tuning changes between task periods (Fig. 2), we fit cosine tuning curves for each neural unit using ordinary least squares regression:

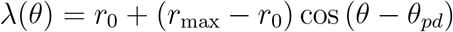

where λ(*θ*) is the estimated firing rate for a given cursor-target direction *θ*. The parameters *θ*_*pd*_, *r*_0_ and *r*_max_ can be interpreted as the preferred direction, the average firing rate, and the tuning amplitude of the unit, respectively. For each neural unit, we fit a separate tuning curve for each task period of the experiment.

We compared the preferred direction *θ*_*pd*_ for each neural unit between Tasks A1 and A2 by computing the average absolute change in preferred direction (Fig. 2b). To calculate the control distribution, for each neural unit, we randomly permuted the task labels for each timestep during Task A1 and Task A2. The difference in preferred direction between Task A1 and A2 was then recalculated using these new task labels.

To visualize how neural activity changes in the 10D population space, we applied linear discriminant analysis to **z**_*t*_, taken in 45ms timesteps, in to order to find the 2D plane that best separates the activity from the three task periods (Fig. 2c). We applied a QR decomposition in order to orthonormalize the basis vectors found by LDA, then projected the neural activity onto this orthonormal basis.

To quantify the changes in population activity between Task A1 and Task A2, we calculated the Mahalanobis distance on a per-target basis between the population activity means across **z**_*t*_, taken in 45ms timesteps, for each task period (Fig. 2d). This distance was computed in the 10D space, using the covariance of the Task A1 neural activity for that target. To calculate the control distribution, for each target, we randomly permuted the task labels for each timestep during Task A1 and Task A2. The Mahalanobis distance between the mean activity for each target was recalculated using the new task labels.

#### Defining the memory trace

Progress quantifies the appropriateness of a particular population activity pattern for a particular BCI map, i.e., the extent to which that population activity pattern drives the cursor towards the target, and is computed as follows. First, we determine the neural push of this activity pattern, **z**_*t*_, through a particular map, *M*_2_, as *M*_2_**z**_*t*_. In equation 4, **m**_0_ and *M*_1_ do not rely on the instantaneous neural activity, and so we do not consider the contributions from these terms. Next, we compute the component of this neural push in the direction of the target. More specifically, for each timestep *t*, we define a unit vector, **c**_*t*_ ∈ ℝ^2×1^, pointing from the current location of the cursor to the target. Thus, the progress at timestep *t* is evaluated as:

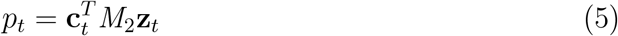

We sought to determine how much more appropriate neural activity is for Map B during Task A2 than it is during Task A1. We call this change in appropriateness a “memory trace” because it measures the lasting alteration of neural activity used during a familiar task (Map A) after a learning experience (Map B). Specifically, we define the memory trace as the difference in progress when neural activity is passed through Map B during Task A2 minus that during Task A1.

#### Testing how Task A2 duration affects the memory trace

We sought to determine whether the memory trace persisted over time (Fig. 4a and Fig. 4b). For each monkey, we divided the sessions into two groups based on whether the Task A2 period was longer or shorter than the median length across all sessions (300 trials). This resulted in 22 sessions in the long Task A2 group (14/22 sessions from Monkey J, average length of 464.36 ± 119.76 Task A2 trials; 8/12 sessions from Monkey N, 400.00 ± 53.45, 0/9 sessions from Monkey L; all values are mean +/-s.d.) and 21 sessions in the short Task A2 group. In order to focus on trials where the monkey had longer exposure to Task A2, we excluded the first 200 trials when calculating the memory trace, leaving at least 100 trials of Task A2 for analysis. For the short Task A2 group, we excluded the first 50 trials of Task A2 as usual (see *Selecting experiments and trials for analysis*).

#### Testing how Task A2 behavior affects the memory trace

We additionally sought to determine whether the memory trace differed as a function of performance through Map A (Fig. 4c and Fig. 4d). To address this, for each target we compared the average progress through Map A during Task A2 to that during Task A1. Targets with acquisition times during Task A2 that were at least as good as Task A1 were placed in the “better behavior group”. There were 21 targets in this group, with an average of 75.0 ± 57.7 ms (mean ± s.d.) faster target acquisition in Task A2 relative to Task A1. Targets which had acquisition times during Task A2 that were worse than Task A1 were placed in the “worse behavior group”. There were 145 targets in this group, with an average of 241.3 ms ± 210.2 ms (mean ± s.d.) slower target acquisition in Task A2 relative to Task A1.

#### Decomposing the memory trace into output-potent and output-null components

In order to determine how the memory trace can coexist without degrading behavioral performance during Task A2, we wanted to determine how changes in neural activity between Task A1 and Task A2 relate to Map A. To address this question, we decomposed neural activity into a component that is output-potent to Map A and a component that is output-null to Map A (Fig. 5). This decomposition was done by applying the singular value decomposition (SVD) to Map A:

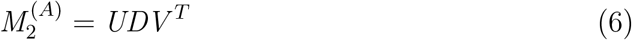

where *U* ∈ ℝ^2×10^, *D* ∈ ℝ^10×10^, and *V* ∈ ℝ^10×10^. *D* is a diagonal matrix, whose diagonal elements are the singular values of 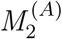. As 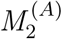 is a matrix of rank two, only the first two diagonal entries of *D* are non-zero. This means that the first two columns of *V* form an orthonormal basis for the output-potent space of 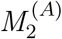. We denote this basis as *R* ∈ ℝ^10×2^. The last 8 columns of *V* form an orthonormal basis of the output-null space of 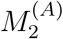. We denote this basis as *N* ∈ ℝ^10×8^.

We can find the component of neural activity potent to Map A as 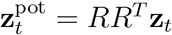. Similarly, the null component is found as 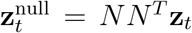. Both 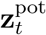 and 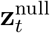 are 10 × 1 vectors, and have the property that 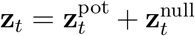. We calculate the potent and null component of the memory trace as before, except utilizing 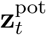 and 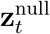 for **z**_*t*_ respectively in equation (4). This decomposition is utilized in Fig. 5 and Extended Data Fig. 5. Note that this decomposition is performed with respect to Map A and not with respect to Map B. This is because, by definition, the memory trace must be in output-potent dimensions of Map B, as those are the only dimensions that determine the cursor velocity through Map B.

#### Path of learning and washout

To distinguish whether the path of washout retraces the path of learning (Fig. 6), we first define the path of learning as the vector in 10D neural activity space from the mean activity during Task A1 to the mean activity during the late Task B period (see *Selecting experiments and trials for analysis*). We similarly define the path of washout as the 10D vector between the mean neural activity during late Task B and the mean activity during Task A2. We then compared the paths of learning and washout by finding the the angle between these two vectors. To obtain a control distribution, for each target, we randomly permuted the task labels for each timestep during Task A1 and Task A2. This mimics a situation in which Task A1 and Task A2 activity patterns come from the same distribution. As task labels for Task B were not shuffled, the paths of learning and washout would thus be equal and opposite on average under this construction. The angle between the paths for each target was recalculated using the new task labels.

### Statistics

Unless otherwise noted, to test for statistical significance, we used nonparametric tests (for example, Wilcoxon signed-rank test or ranked-sum test), which do not assume normality. All P-values less than 10^−10^ were reported as *P <* 10^−10^, regardless of how small the P-value was.

### Data availability

The data that support the findings of this study are available from the authors upon reasonable request.

### Code availability

Python code that supports the data analyses will be made publicly available upon publication.

**Extended Data Fig. 1.**
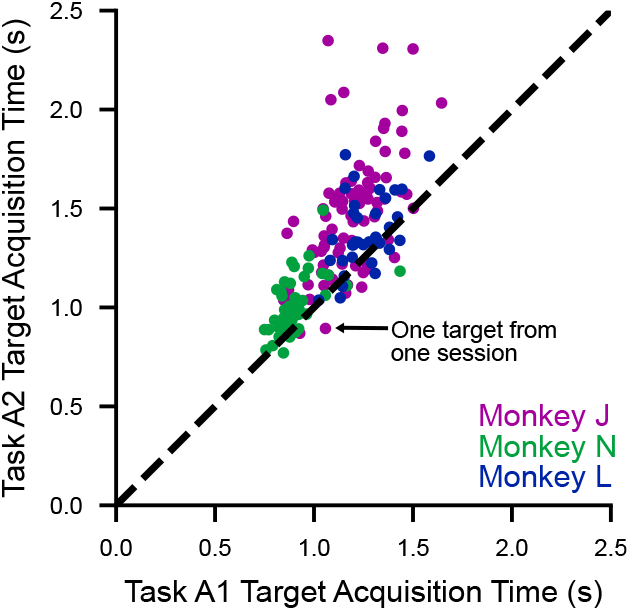
Comparison of behavioral performance in Tasks A1 and A2. Here we plot the average acquisition time for a given target during Task A1 against its average acquisition time during Task A2. Performance in Task A2 tended to be lower than performance in Task A1, likely due to satiation or fatigue. In Fig. 4 and Extended Data Fig. 3 we demonstrate that this difference in behavior is not the cause of the memory trace. The targets that fall below the diagonal are those in which performance during Task A2 is better than during Task A1, and are the same targets that are included in the “better behavior group” in Fig. 4d.

**Extended Data Fig. 2.**
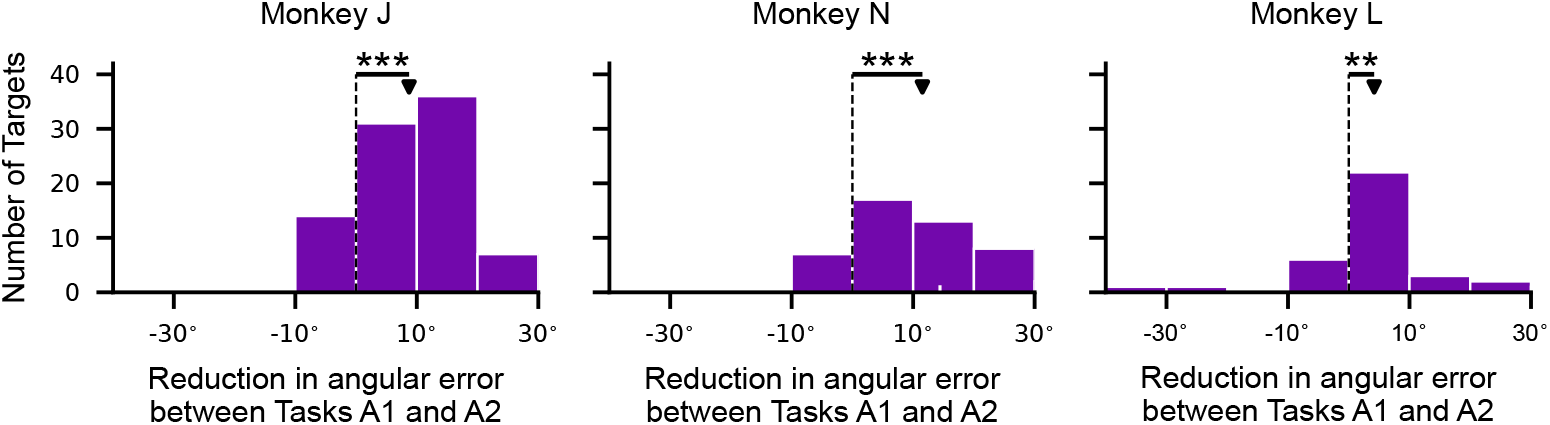
A memory trace is also evident when measured using angular error. In Fig. 3d, we measured the memory trace using “progress”, which is defined as the velocity by which a neural activity pattern would have moved the cursor toward the target (see Methods). We could have alternatively measured the memory trace in terms of angular error instead of progress. In contrast to progress which depends on velocity magnitude and direction, angular error depends only on the velocity direction. Angular error is defined at each timestep as the angular difference between the velocity vector of the neural push and the cursor-to-target direction. As with progress, the velocity of the neural push is defined using the Task A1 (or Task A2) neural activity projected through Map B. We then compute the angular error for Task A1 minus the angular error for Task A2. We use unsigned angular error so clockwise and counterclockwise errors do not cancel each other out when averaging. Smaller angular errors are better. Thus, when angular error is smaller for Task A2 relative to Task A1, a memory trace is present (*P <* 10^−10^, two-sided paired Wilcoxon sign-rank test, n=88 targets; Monkey N, *P* = 1.02 × 10^−7^, n=48; *P* = 0.0027, n=36 targets). The white tick mark on the horizontal axis of the middle histogram denotes the example target illustrated in Fig. 3a, Fig. 3b and Fig. 3c.

**Extended Data Fig. 3.**
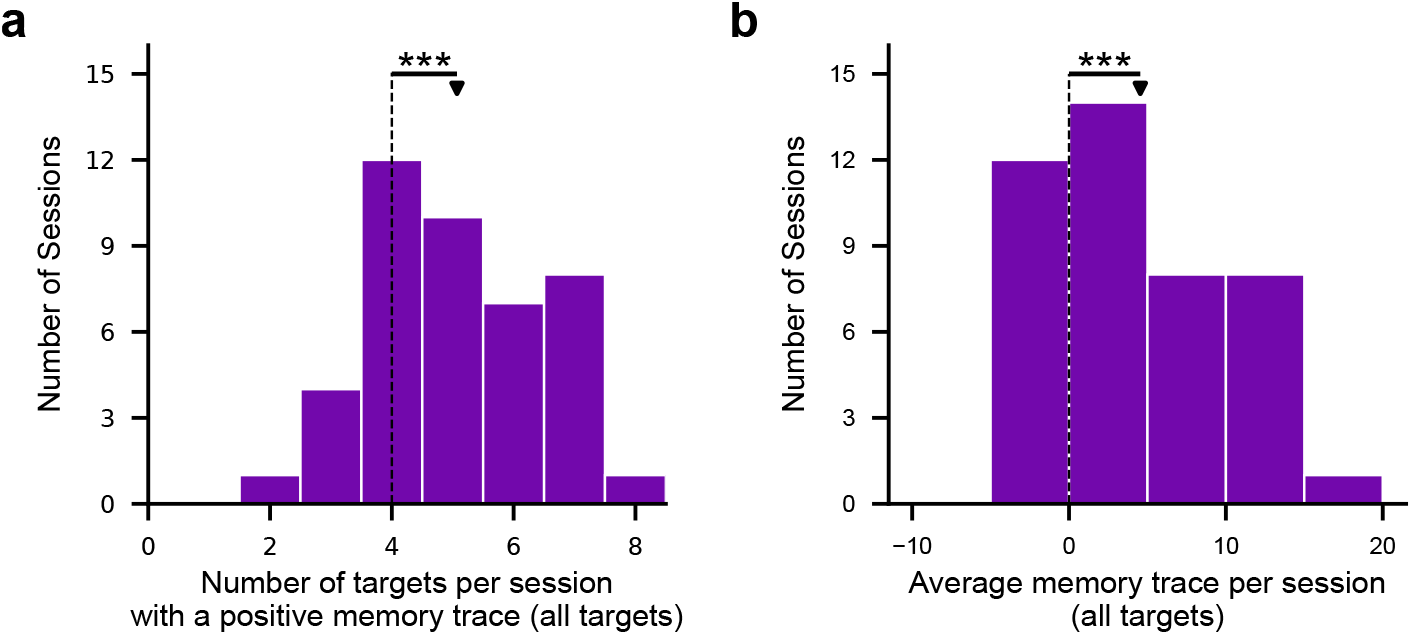
Most targets per session exhibit a memory trace. We considered whether the memory trace could be due to a global shift in neural activity (e.g., due to a neural recording instability) that leads to an increase in Map B progress for some targets at the expense of progress for targets on the opposite side of the monkey’s workspace. If this were the case, we would expect that only half of the targets in each session, including targets that show little or no learning, would show a memory trace. **(a)** Instead, we found that more than half of the targets per session showed a memory trace (*P* = 7.21 × 10^−5^, two-sided paired Wilcoxon signed-rank test, n=43 sessions across monkeys). **(b)** Similarly, we found that the average memory trace across all eight targets per session is positive (*P* = 4.25 × 10^−5^, two-sided paired Wilcoxon signed-rank test, n=43 sessions across monkeys).

**Extended Data Fig. 4.**
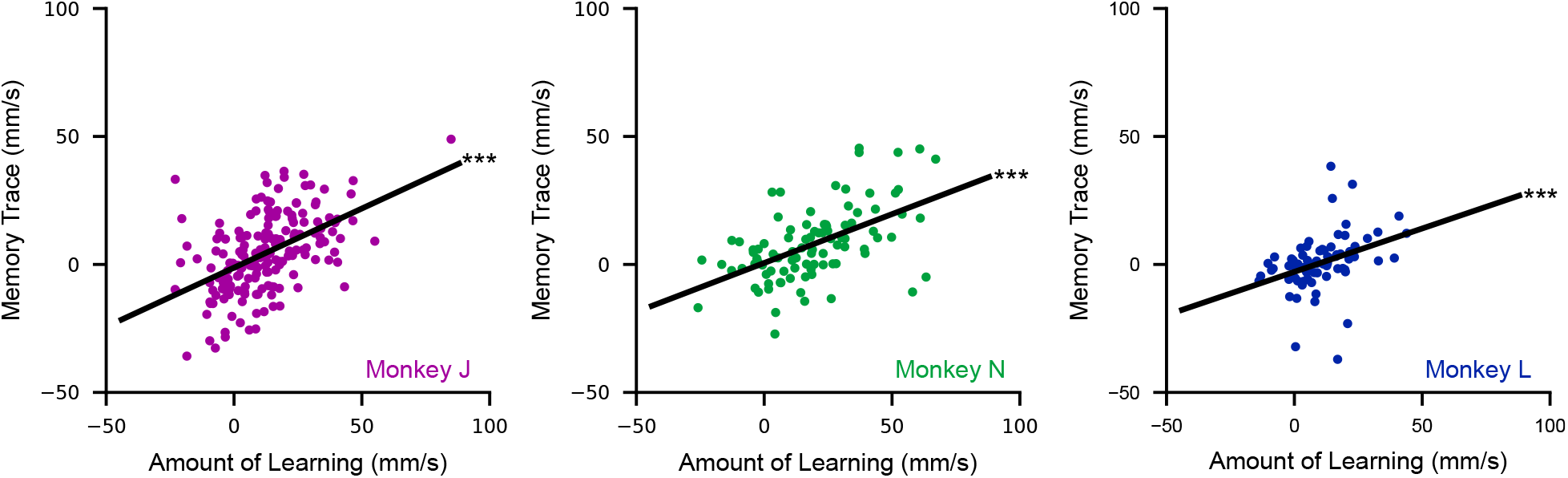
The amount of learning is correlated with the size of the memory trace. If the memory trace arose spuriously and not as the result of the learning experience, we would expect the size of the memory trace (measured using neural activity during Task A2) to be uncorrelated with the amount of learning (measured using neural activity during Task B). The amount of learning during the Task B period positively correlates with the magnitude of the memory trace (Monkey J *R*^2^ = 0.25, *P <* 10^−10^, one-sided F test, n=176 targets; Monkey N, *R*^2^ = 0.29, *P* = 1.23 × 10^−8^, *n* = 96; Monkey L, *R*^2^ = 0.13, *P* = 0.0017, *n* = 72). All targets were included in this analysis, though similar results hold when only examining well-learned targets. We considered the possibility that these results could have arisen trivially due to the memory trace and amount of learning both being calculated relative to Map B progress during Task A1. We thus reran this analysis without subtracting this quantity (that is, regressing Map B progress during Task B with Map B progress during Task A2) and arrived at similar results (Monkey J *R*^2^ = 0.48, *P <* 10^−10^; Monkey N, *R*^2^ = 0.45, *P <* 10^−10^; Monkey L, *R*^2^ = 0.37, *P* = 1.14 × 10^−8^). This supports the notion that the memory trace is the result of the preceding learning experience. Furthermore, in Fig. 3d, Monkey L showed a smaller memory trace than Monkeys J and N. While the scatter of values for Monkey L lies within the scatter of Monkeys N and J, Monkey L showed less learning on average than Monkeys J and N. This is a possible explanation for Monkey L’s smaller memory trace.

**Extended Data Fig. 5.**
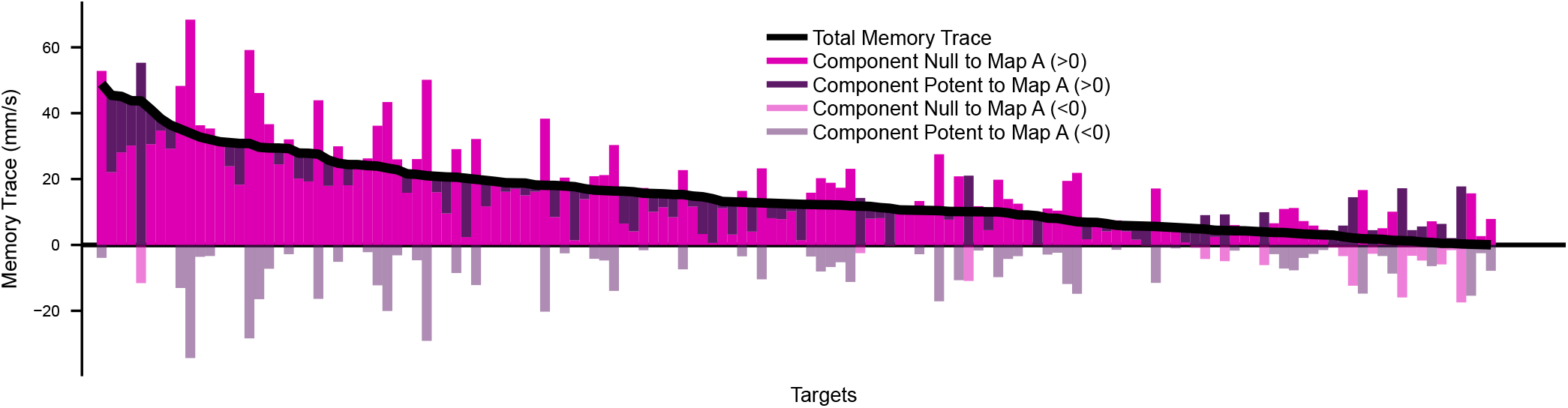
The majority of the memory trace resides in dimensions outputnull to Map A. To understand which dimensions of neural activity contribute to the memory trace, we decomposed neural activity into components that are output-potent and output-null to Map A and evaluated their contribution to the memory trace (Fig. 5). Here, we breakdown Fig. 5 by target. Targets across all sessions and monkeys are ordered by the total memory trace expressed for that target (black line). The contributions by the potent and null spaces of Map A are shown in purple and magenta, respectively. As the total memory trace is the sum of the contributions from the output-potent and output-null components, it is possible for one of these components to have a negative contribution and the total memory trace to still be positive. A negative value indicates progress through Map B is smaller during Task A2 relative to Task A1 for that component. For visual clarity, we use dark shading for positive values and light shading for negative values. For a given target, there is one purple bar (light or dark) and one magenta bar (light or dark). We find the majority of the memory trace lies in resides in dimensions output-null to Map A (magenta bars tend to be larger than purple bars), as quantified in Fig. 5.

